# A chronic state biostimuli-responsive therapeutic model system: applications to skin tissue repair and regeneration

**DOI:** 10.1101/2025.11.24.690306

**Authors:** Tramata Djakite, Aline Échalard, Béatrice Labat, Anthony C. Duncan

## Abstract

We report on the elaboration of a biocompatible therapeutic model system (TMS) capable of detecting the state of chronicity of a wound and responding to the latter by releasing the appropriate healing agent (antibiotic but not only). Different formulations of PVA-borate hydrogel systems containing methylene blue (MB) and levofloxacin (L) drug antibiotic were prepared. We were able to demonstrate sensitivity to microenvironmental factors such as pH, and high levels of ROS – a major biomarker of a chronic inflammation state. Furthermore, these reversible gels “open” and enact an ROS triggered drug delivery response. These results are promising in that they provide a potent wound dressing layer capable of sensing the chronic state of a wound and responding conditionally to the latter by delivering the appropriate amount of drug only when required. Furthermore, our system may be optimized for chronic wound dressing healing and promote better wound care by limiting unnecessary overexposure of the patient to the antibiotic. Our system provides a potent novel tool to the arsenal of existing strategies aiming at combatting bacterial resistance to antibiotics – a global societal challenge. Our results additionally confirmed good biocompatibility for skin tissue regeneration.

## Introduction

Due to their numerous potential applications, stimuli-responsive PVA hydrogels have gained increasing interest in numerous and unexpected fields (1–8). In the biomedical area these include applications such as controlled drug release devices, skin tissue engineering and tissue regeneration systems, chronic wound care to name a few. They also present a real benefit in the context of chronic wound treatment.

Chronic wound (CW) care is a serious societal and human suffering problem affecting 1-2% of world population and costing up to nearly $150 billion according to some estimates (9–12). Wound healing is a complex, dynamic, time-dependent multivariable process. Conventional “passive” wound dressings exhibit a few disadvantages, one being their inability to respond adequately to the rapidly evolving wound microenvironment for optimal healing (13). Chronic wounds are wounds that stay blocked in the inflammation step of the healing process due notably to the presence of bacterial biofilms. In previous work we were able to bring solutions to overcome this impediment by the development of multiple layer multifunctional wound dressings – the external layer protecting against external infections, the internal layers presenting the ability to release bioactive agents to promote a microenvironment that favors or hastens wound healing (14–16). In that respect, the latter function may be perfected or rendered more efficient in specific required situations via the use of tailored biofunctional hydrogels. In point of fact, these emerge as a promising strategy due to their ability to provide adaptive microenvironments and appropriate responses in a timely fashion. Furthermore, due to their adequate intrinsic properties, they are potent candidates for drug release, tissue repair/regeneration, tissue engineering and wound dressing applications. Biostimuli-responsive gels, in particular, have gained particular attention for their promising potential in the context of wound healing and skin tissue regeneration [(17), (18), (19), (20), (21)].

PVA functional hydrogels are attractive for skin wound healing and regeneration due to their inherent properties: water absorbing capacities (e.g. wound exudate absorbing capacity), ability to be irreversibly or reversibly crosslinked and functionalized in many ways due to their abundant backbone chain hydroxyl groups. These may also be readily tailored to the required specifications for a particular wound dressing application (e.g. appropriate internal moisture; adequate O2, CO2 gas and water vapor exchange permeability; protection from external bacterial contaminations) as we described previously (14–16).

Furthermore, growing bacterial resistance to formerly efficient antibiotics is a major societal problem (10). One main reason for this is their overuse in situations that do not require their high-level involvement.

In the context of chronic wound treatment, the elaboration of release systems that liberate the antibiotic (or antibiofilm) drug in appropriate amounts, solely when required emerges as a promising tool.

In this paper we explore the elaboration of a bio-responsive cost-effective controlled release system capable of achieving the aforementioned objectives. The system has the ability of releasing the appropriate drug solely in response to the detection of high levels of a specific biomarker of a chronic state – reactive oxidative species (ROS). Although in this article we treat the potential chronic wound healing aspect of our system, it is not excluded that it may also exhibit non negligeable potential in other situations where ROS is also involved as a biomarker of a pathological state – cancer for instance (22–24).

## Materials and methods

PVA (98-99% hydrolyzed, Mw 88-98 kg.mol^-1^) was purchased at Alfa Aesar. Anydrous sodium tetraborate, Borax, 381,37 g/mol 99% pure) was purchased at Prolabo. Methylene blue (MB, 95% pur) was purchased at Accroc Organics. Levofloxacin (HPLC grade) was purchased at Sigma Aldrich. Hydrogen peroxide (60% w/v) was purchased at Fisher scientific.

## Experimental

### A PVA based biostimuli-responsive therapeutic model system (TMS)

Three families of physical PVA gels called TMS, TMS (+) and TMS(L+) were elaborated. The description for each gel is found in table 1.

**Table 1.** Nomenclature and description of the prepared hydrogels.

| Nomenclature | Description |
| --- | --- |
| TMS | PVA 3% (w/w) - borax 6% (w/w) |
| TMS (+) | TMS prepared in a solution of MB (8 µg/ml) |
| TMS (L+) | Gel TMS prepared in a solution of levofloxacin (30 µg/ml) |

The gels were prepared as follows.

### TMS gel elaboration

<u>Procedure 1</u>: firstly, two solutions A and B were prepared. *Solution A* was a 3% w/w PVA aqueous solution obtained by dissolving 3 g of PVA into 97 g of pure deionized water, and heated between 80°C and 90°C using a water bath until total dissolution was achieved and a clear transparent homogeneous solution was obtained (no crystal remaining). *Solution B* was a 6% w/w borax solution obtained by dissolving 6 g of the solute into 94 g of deionized water at a steady temperature of 30 °C in a water bath. Both solutions A and B were allowed to cool down to room temperature. Then, 10 ml of solution A was progressively adding dropwise to 10 ml of solution B, while slowly stirred with magnetic bar at room temperature, until a final uniform gel was obtained (TMS).

### TMS(+) gel elaboration

Preparation of the methylene blue (MB) containing gel TMS(+) was obtained as follows. Methylene blue (MB) was dissolved in pure water to obtain via serial dilutions a final concentration of 8 µg/ml (solution C). Then, a solution D was elaborated by preparing a 3% w/w PVA in solution C using the same dissolution process as described for solution A. TMS(+) was finally obtained by progressively adding dropwise 10 ml of solution B into 10 ml of solution D, while slowly magnetically stirring until a final uniform gel was obtained.

### TMS(L+) gel elaboration

Preparation of TMS(L+) was obtained as follows. Levofloxacin (L) was dissolved in pure water to obtain a final concentration of 30 µg/ml (solution E). 3% w/w PVA was prepared in solution E according to same process as described for solution A to obtain a final solution F. Then, 10 ml of solution B were progressively added dropwise into 10 ml of solution F, while slowly magnetically stirring at room temperature, until a final uniform gel was obtained (TMS(L+)).

### Characterization

#### FTIR

After previously being air dried at ambient temperature for 24 hours, the gels were analyzed using an FTIR-ATR Bruker Tensor 27 spectrometer. All spectra were performed in the 600-4000 cm^-1^ range on germanium crystal with an incident beam of 60°.

#### UV-Vis spectroscopy

Drug release from gels were analyzed using the Spectronic Unicam UV 300. Maximum absorbance wavelengths (max) were determined by scanning at wavelengths between 200 and 800 nm. These were found to be 626 nm and 259 nm for the MB and levofloxacin, respectively. Therefore, absorbance readings were performed at those maximal wavelengths for the latter.

#### Stimuli responsiveness

Gel responsiveness to temperature, pH and ROS agents, were evaluated.

##### Temperature dependent response

0,5 g of the gel were placed in an Eppendorf tube and 1 ml of deionized water was added. The whole was immersed in a water bath at low (0°C) than high temperatures (50°C) during 30 min. This warm/cold cycle was performed 5 times in a row.

##### pH dependent response

In order to evaluate TMS pH dependence response, 1 ml of a predetermined pH value solution was added to 0.5 g of the gel. The acidic versus alkaline test environments were HCl and NaOH aqueous solutions. Exposure times to the latter were 1,2,3 and 24 hours. Occurrence of gel-sol transition (GST) in such conditions were monitored visually. The transition was confirmed by turning the Eppendorf tube up-side down. Gel-sol transition occurred if the tube content flowed down the sides of the Eppendorf tube to the bottom. No sol-gel transition was witnessed by the gel staying stuck up in the top of the tube.

##### ROS dependent response

Hydrogen peroxide (H2O2) was used as a model reactive oxidative species (ROS) molecule. To evaluate ROS sensitivity of our gels, H2O2 aqueous solutions at various concentrations was prepared via sequential dilution from an initial 60 % w/w hydrogen peroxide commercial solution. In an Eppendorf tube, 1 ml of hydrogen peroxide at a given concentration was added to 0,5 g of the gel. After a given exposure time (30 minutes) gel-sol transition was monitored. Exploring a wide range of concentrations between 17.65 M (60 %) and to 0.001mM, allowed us to determine a Minimal Transition Concentration (MTS) ROS sensitivity range for the studied gel.

### Biocompatibility evaluation

#### Cell culture

L929 mouse fibroblast cells were used to qualitatively and quantitatively assess cell viability. They were routinely grown into 75 cm^2^ plastic tissue culture flasks, fed with complete culture medium (DMEM culture medium supplemented with 10% fetal bovine serum (FBS), streptomycin/penicillin (100 mg mL^-1^/100 mM respectively) and 2 mM L-glutamine) and incubated in a controlled humidified atmosphere with 5% CO2 at 37°C. Then subconfluent cells were enzymatically detached using 0.25% trypsin–1 mM EDTA (Sigma) and seeded at a density of 20 000 cells.cm^-2^ either within 12-wells plates (for optical microscopy observations and the Alamar Blue® assay), or within 12-wells plates with glass slides at their bottom (for the Live/Dead® assay).

#### Indirect cytotoxicity tests

Cytotoxicity of all the hydrogel formulations was assessed through an eluting test method, according the standard ISO10993-5 (25). Two series of TMS (L+) and TMS hydrogels, respectively, were prepared. The first one consisted of TMS (L+) and TMS that were air-dried for 24h (t24) and the second one used immediately after elaboration (t0). All the samples were UV-sterilized (30 min, λ = 254 nm). Then, to collect any potentially cytotoxic compounds released from the hydrogels, in Eppendorf tubes they were exposed for 24 h at 37°C to complete culture medium, at a ratio of 0.1g hydrogel per mL and kept as such in sterile conditions until used. The solutions were named as “extracts”.

Cell metabolic activity was finally assessed with Alamar Blue® assay (AB – Bio-Rad, UK), 24h after cells contacting the extracts and the controls. AB is commonly used to quantitatively assess cell metabolic activity since it is based on oxidation-reduction reactions occurring in cell mitochondria. For that, 10% (v/v) AB-containing standard culture medium was given to cells for 3h. Then, absorbance measurements were performed at 570 and 600 nm with a microplate reader (BioTeck – Synergy 2, USA).

#### Direct cytotoxicity tests

After 24 h of standing in the extracts, cells were observed with the use of an inverted optical microscope (Motic, Spain) equipped with phase contrast illumination and a CCD camera (Zeiss) to inspect their global morphology. They were compared to negative and positive cytotoxic controls, which corresponded to cells cultured in standard culture medium alone or supplemented with 10% DMSO.

#### Live/Dead ® cell tests

In parallel, Live/Dead® assays were performed to qualitatively assess cell viability. For that, the cells were seeded at the bottom of 24-well plates on top of sterilized glass covers. The extracts were then distributed over all cells. After 24h, cells were observed under epifluorescence microscopy (Zeiss Axio Scope A1, Carl Zeiss, Germany) equipped with a CCD camera (Zeiss). Cells that appeared in green were viable cells whereas those that appeared in red were considered as damaged or dead cells. Images were post-treated with ImageJ to merge green and red channels.

## Results & discussion

The final gels obtained are shown in Figure 1. They exhibit high malleability and self-repairing properties in the sense that after being broken up into multiple parts, the latter, once brought together in contact again, recombine into the initial cohesive homogeneous gel. The gels also present high water content with a gelatinous aspect. These properties are appreciable for chronic wound dressing or tissue reparation/regeneration applications since for these reasons the gel will tend to mimic the host extracellular matrix of the wound.

**Figure 1.**
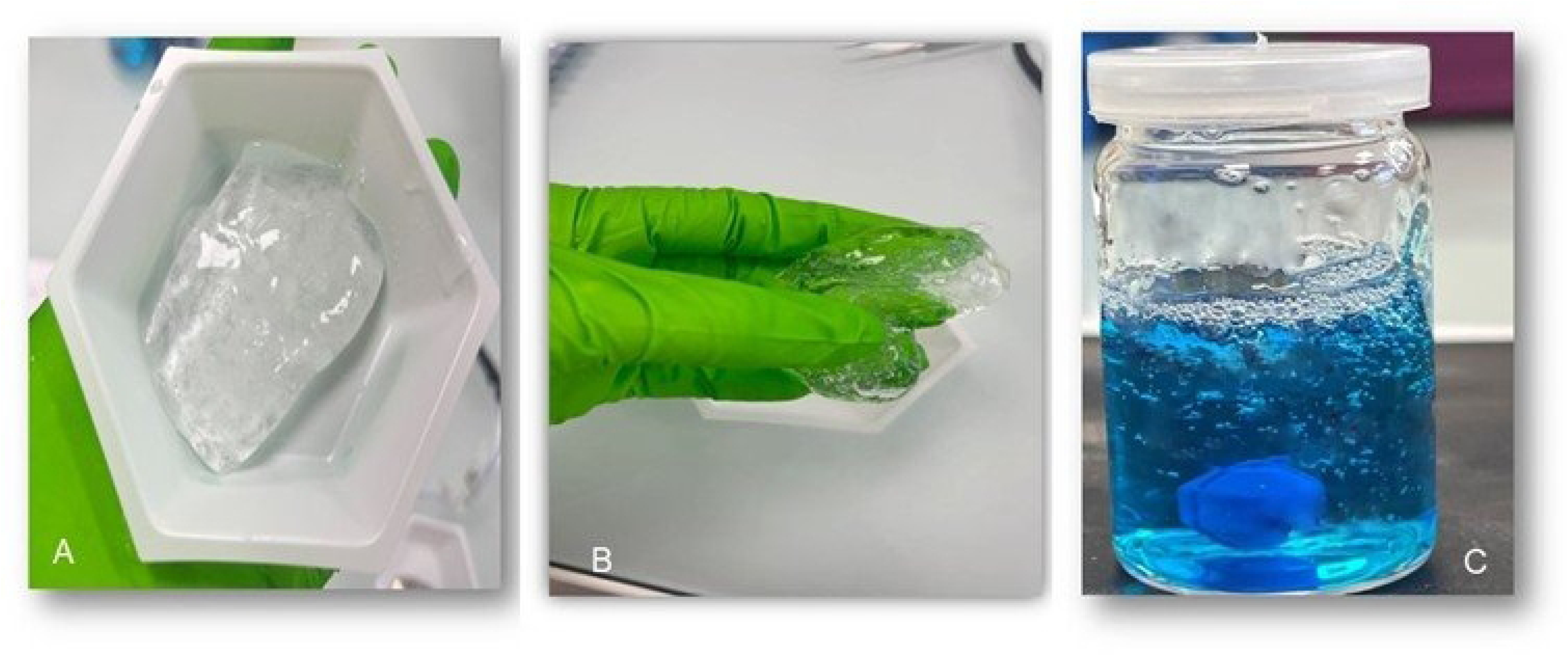
Visual aspects of TMS (A), TMSL (B) and TMS+ (C) gels.

### Temperature response

It is worth noting here that for the gel-sol transition (GST) studies, presented here below, the gel was placed at the bottom of the Eppendorf tube. The latter was then turned upside. If gel-to-solution transition took place the resulting liquid would thus flow down to the bottom by gravity. If no GST occurred, the gel will stay “stuck” in the upper part of the tube. Thi is the case in figure 2, the after thermal cycling treatment of gels performed during 24 h per half cycle between 50°C and 4 °C. The latter treatment had little effect on visual aspect or mechanical integrity of the gel (Figure 2). These results suggest that our gels are stable to temperature fluctuations within the range studied.

**Figure 2.**
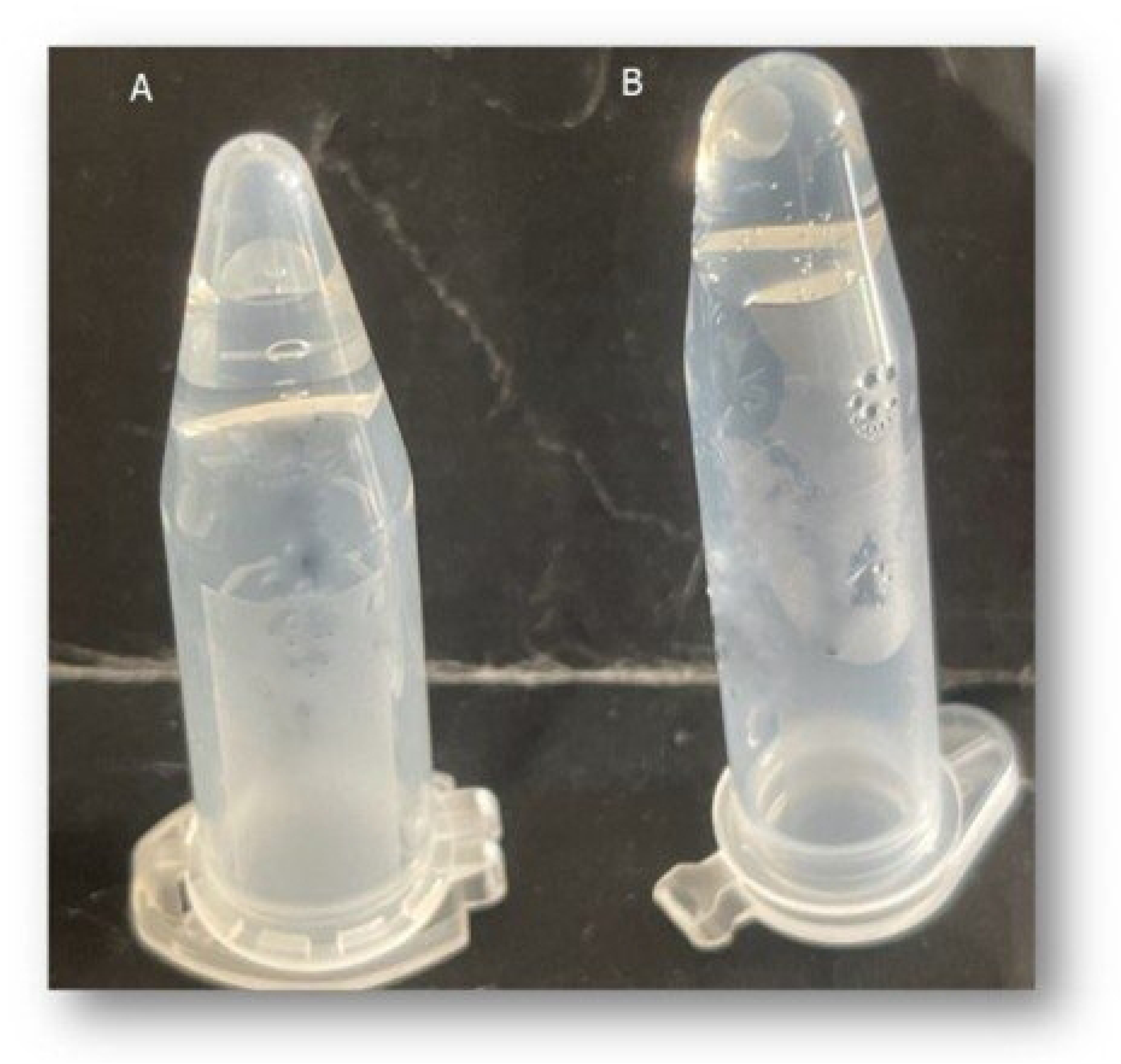
Gel TMS (-) temperature cycling response. Results shown are subsequent to 5 thermal cycles between 50°C and 4°C. (A) TMS (-) control (no cycling). (B) TMS (-) after 5 cycles.

### pH stimulus response

TMS gel response to different pH environments are shown in figure 3. At low and neutral pH values, total and partial GST were observed, respectively. In contrast, no GST was observed at high pH value. Thus, the alkaline environment appeared to have a structuring effect on the gel. Whereas an acidic environment clearly had a destructuring effect, as evidenced by the observed the total GST (Figure 3B). This observation is consistent with the nature of the PVA polymer inter-chain crosslink: a complex favored by the presence of abundant hydroxide ions.

**Figure 3.**
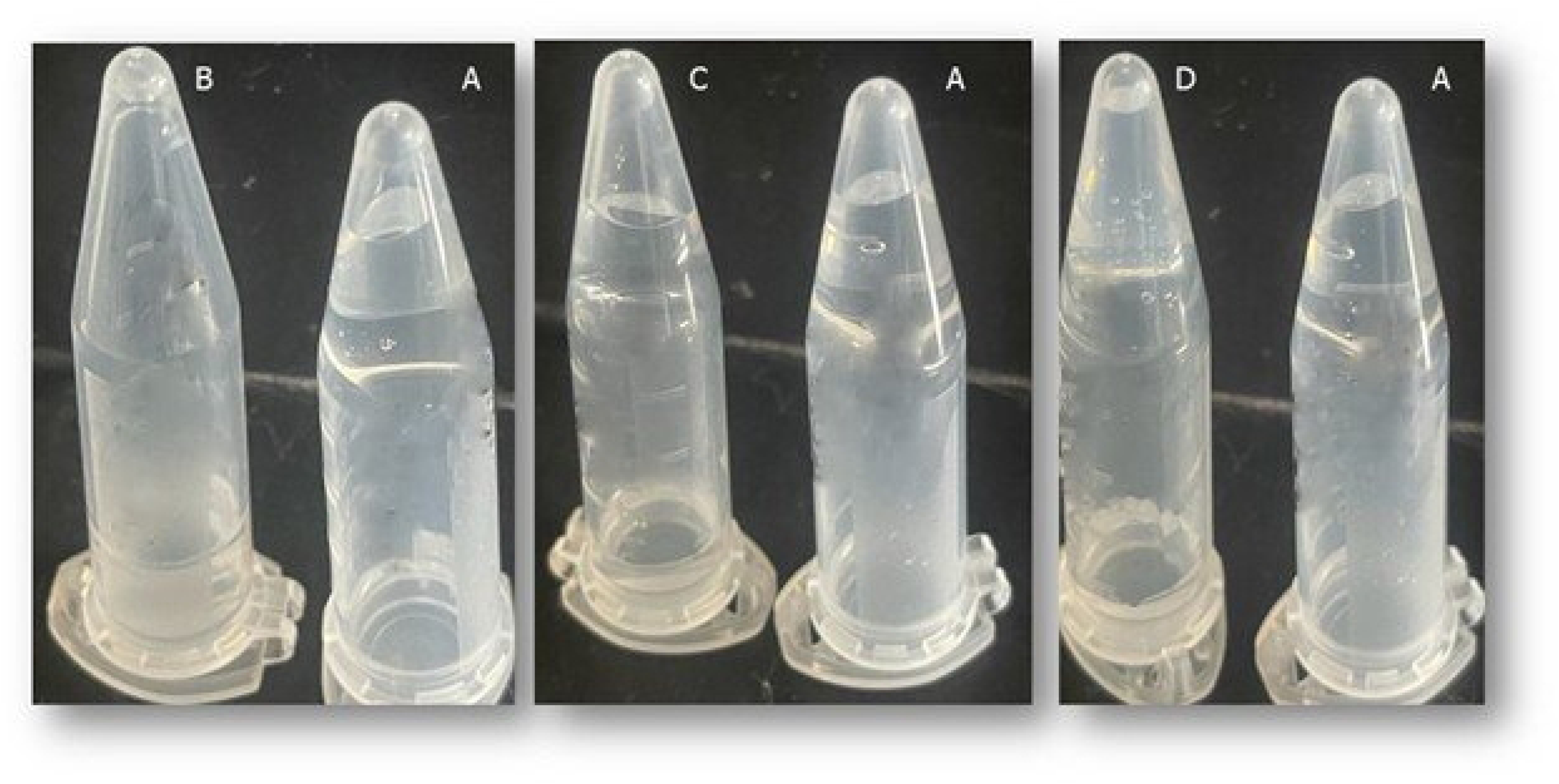
TMS pH dependent response. TMS gel control alone (A) and exposed to different pH values (B) 0.6, (C) 7 and (D) 14 over 30 minutes.

pH response for the MB containing gels (TMS+) is shown in figure 4. Total and partial sol-gel transition were observed at low (figure 4B) and intermediate pH values (figure 4C), respectively. TMS+ gels remained solid in alkaline conditions (figure 4D). These MB containing gels TMS+ results are akin to those observed for the TMS gels (without MB). They suggest little effect (little interference) of the presence of the methylene blue on the PVA inter-chain crosslinking occurence.

**Figure 4.**
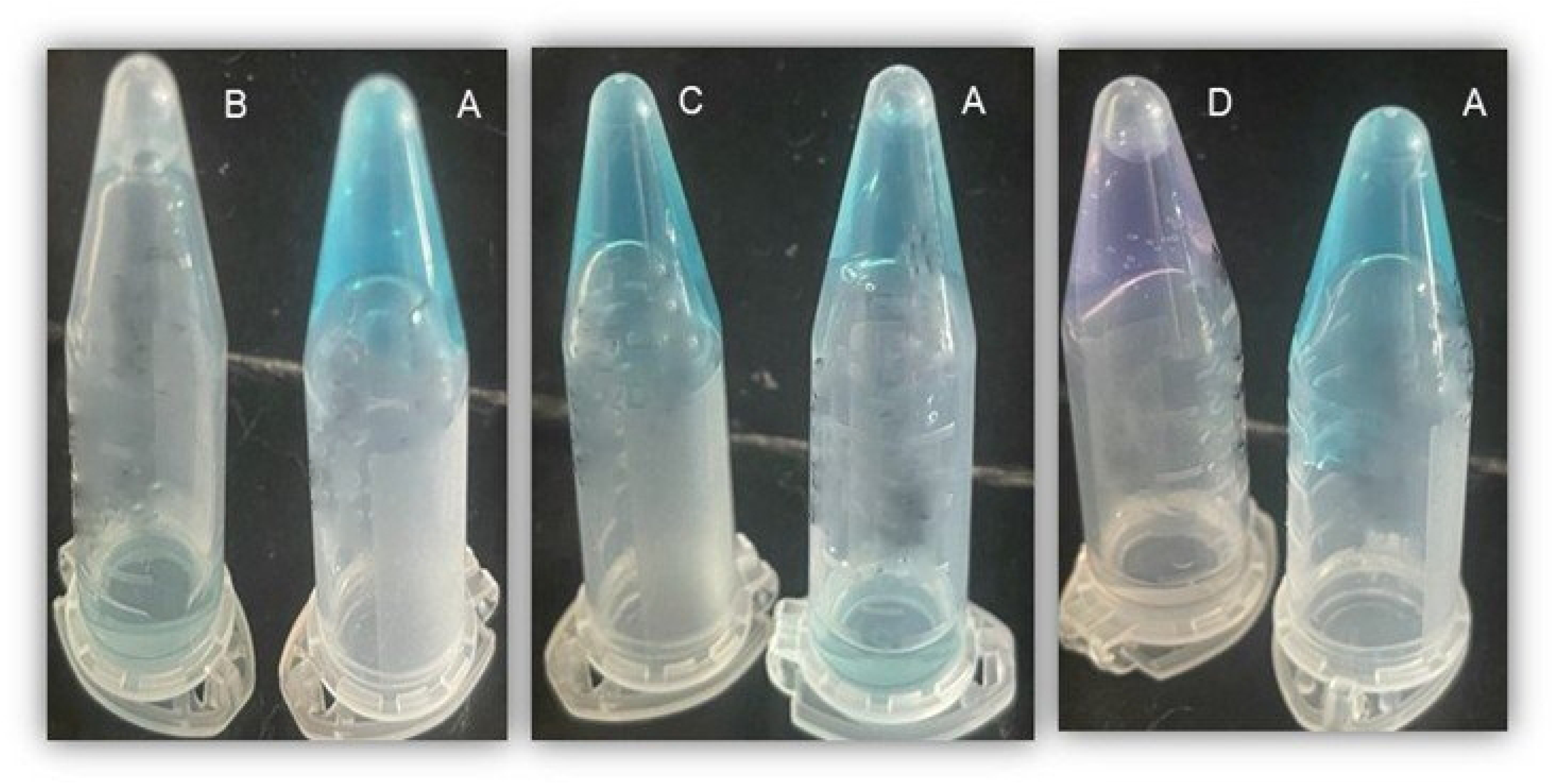
TMS+ pH dependent response. TMS(+) gel control alone (A) and exposed to aqueous solution at the following pH values (B) 0.6, (C) 7 and (D) 14.

It is worth noting here that the TMS+ gel shows a net color change at high pH value from blue to violet (Figure 4D). A possible explanation for this is the chemical reactions associated with OH^-^ ions and MB in alkaline environments, yielding a family of dye (by)products such as methylene azure (azure B) and methylene violet as previously described (26) (27).

Results for the levofloxacin containing gels (TMSL) are shown in figure 5. These slightly differ compared to those for the TMS+ gels. Indeed, GST was observed not only at low (Figure 5B) but also at intermediate pH values (Figure 5C). More surprisingly, partial GST was also observed at high pH value (Figure 5D). The latter suggests that, in contrast to MB, the presence of the levofloxacin drug (L) appears to induce some kind of interference in the PVA inter-chain crosslinking mechanism.

**Figure 5.**
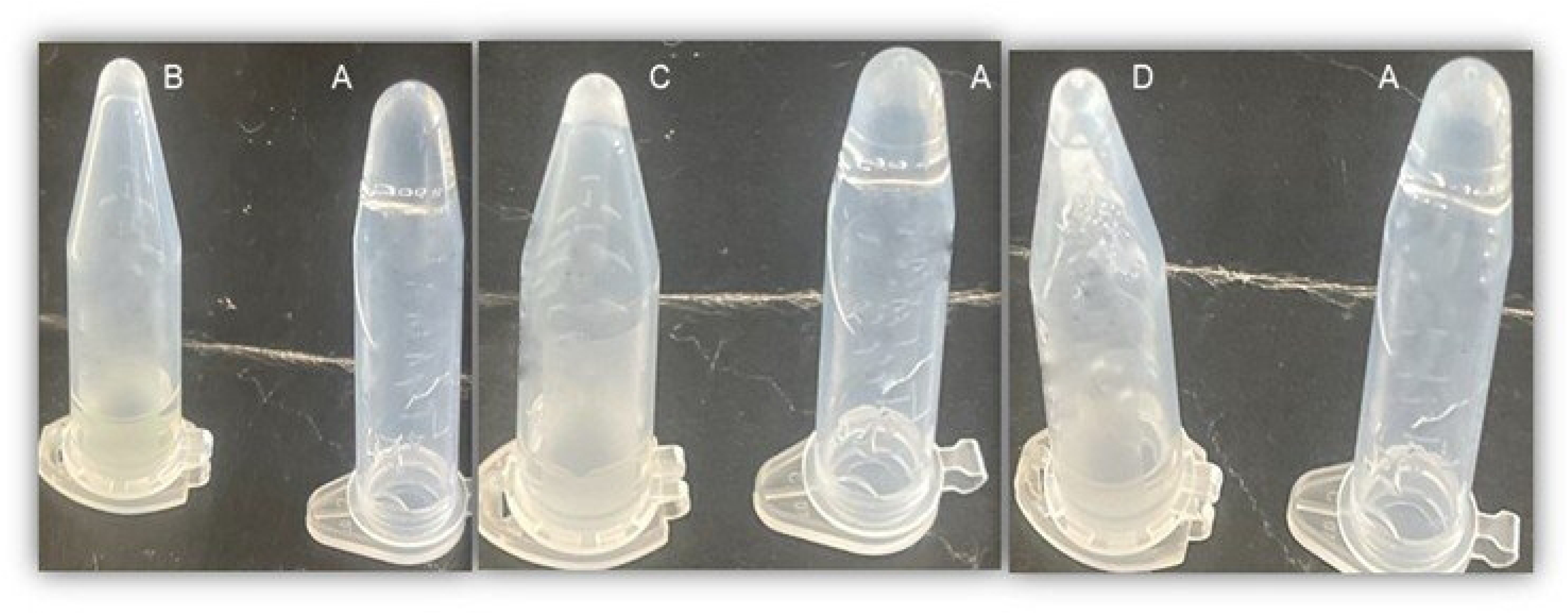
TMSL pH dependent response. TMSL gel control alone (A) and exposed to each aqueous solution at pH value equal to 0.6 (B), 7 (C) 7 and 14 (D).

These observed differences may be due to molecular drug size differences, levofloxacin being a bulkier molecule compared to the more linear MB. Another explanation pertains to pH dependence of levofloxacin net charge. Indeed, MB *net charge* is independent of pH value. In contrast, that of levofloxacin evolves with respect to pH value. This is inherent to the three intrinsic pKa values for levofloxacin (supporting information).

Indeed, according to its pKa values, levofloxacin (L) is expected to have a total negative net charge at high pH values. Thus, electrostatic repulsion is highly probable between the latter (L) and the negatively charged borate PVA-interchain crosslinks in alkaline environment. This would in turn contribute to cohesion weakening observed of the TML gel.

### TMS response to ROS stimuli

Gel exposure to ROS results are shown in figure 6. GST was observed for all gel families TMS, TMS+ and TMSL confirming good ROS sensitivity for all gels.

**Figure 6.**
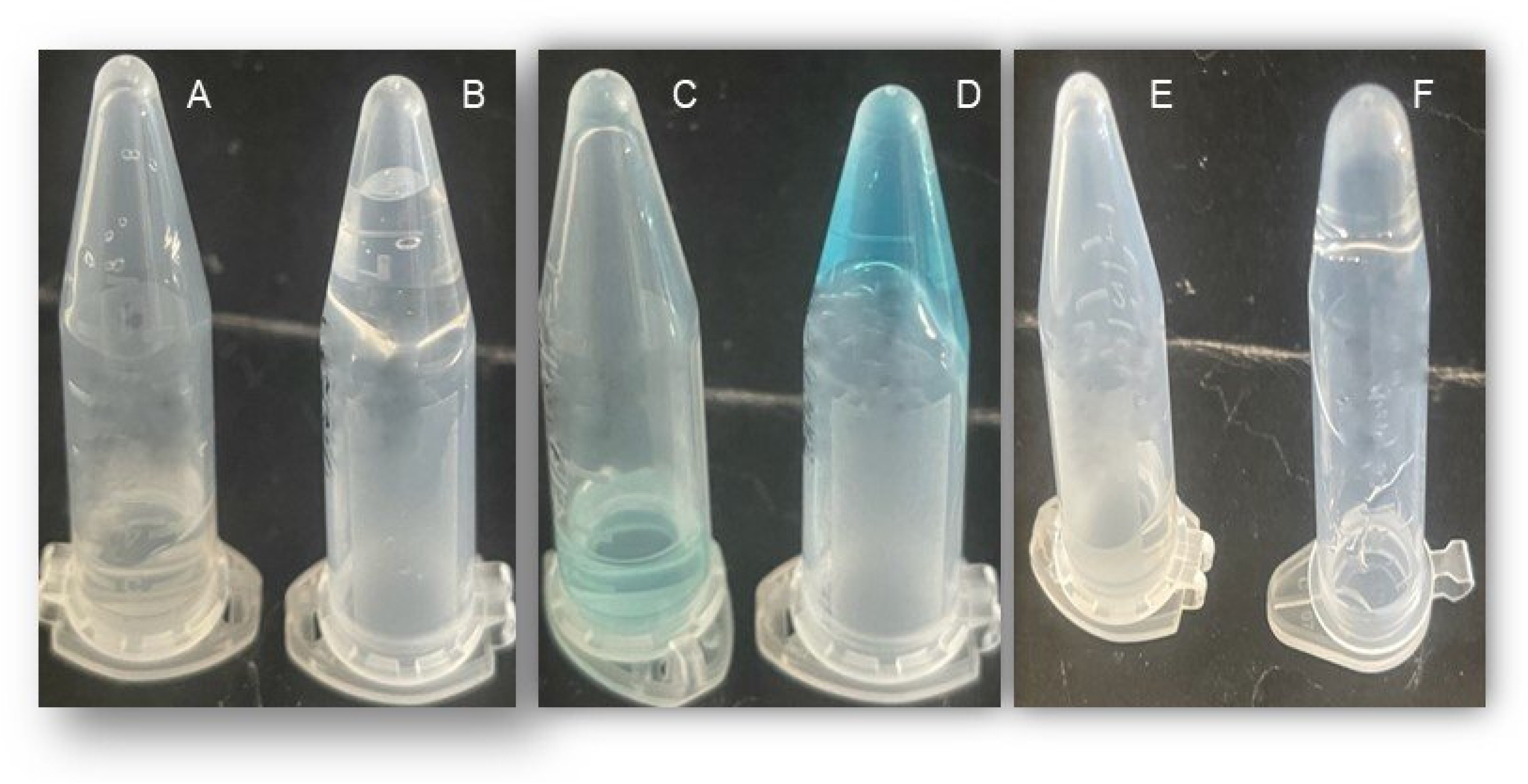
ROS dependent response. TMS (B), TMS+ (D) and TMSL alone (F) and respectively exposed to a 60% w/v hydrogen peroxide solution (A, C, E).

### FTIR analysis

TMS gel spectrum versus standard PVA is shown in supporting information. Characteristic peaks at 3393 (A), 2980 (B), and 1649 cm^-1^ (C) for the abundant hydroxyl groups of PVA, aliphatic and carbon-bore-oxygen stretch (C-O-B), respectively, were clearly visible in the TMS gel spectrum as described previously (5). The results are consistent with the PVA-borate crosslinking within the TMS gels (supporting information).

### Determination of a minimal ROS concentration range (MCR) for GST occurrence

Test results for the determination of minimal concentration range of H2O2 for GST occurrence are shown in the supporting information. GST occurred for concentrations as low as 1 µM H2O2 for both TMS and TML gels (in a proportion of 0.5 g gel/ 1 mL of H2O2 solution). Both latter exhibited GST at 20° and 37°C within 30 minutes.

These results are promising since they suggest that sol-gel transition would likely occur at body temperature for the levofloxacin loaded gel thus liberating the antibiotic in the presence of the ROS chronic state biomarker. As shown aforementioned GST is solely due to the presence of H2O2 and not to temperature influence (since gel remains stable at 37°C and above without ROS present).

### Drug release studies

The impact of environmental pH on drug release from the gels measured over a period of 24 hours is shown in Figure 7.

**Figure 7.**
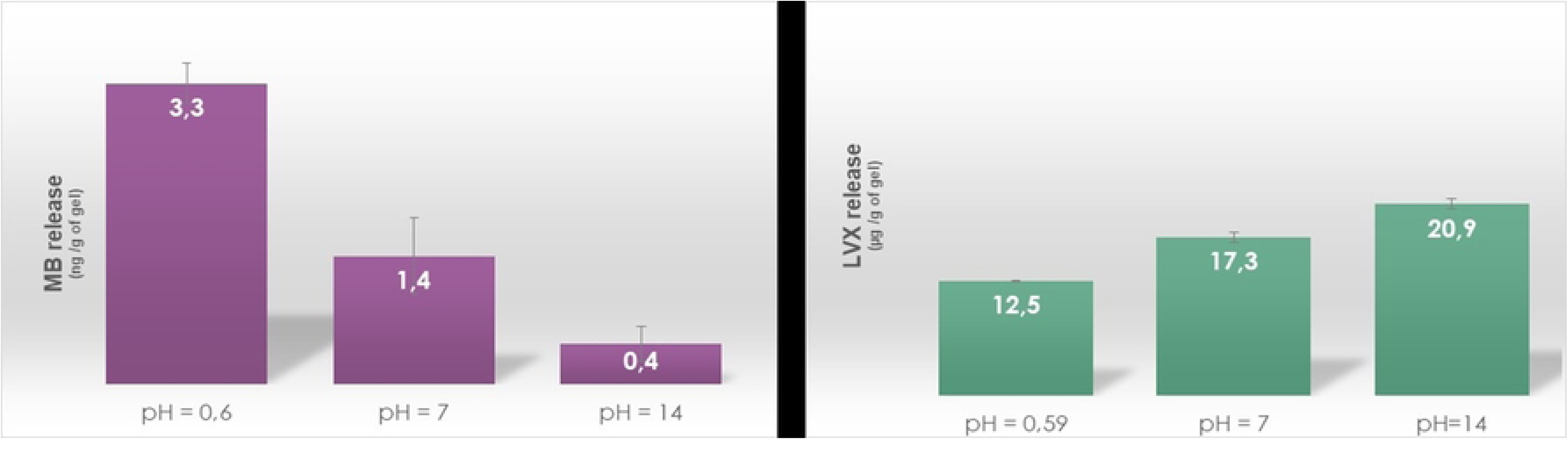
Drug release. Methylene blue (left) and levofloxacin (right) release from TMS(+) gel at acid, neutral and alkaline pH values over a 24h period. Results are shown as average and standard deviation (n = 5).

One first observation is that MB release results are consistent with GST results in an acidic environment, since highest MB liberation rates and total GST were observed at lowest pH value (Figure 7, left). Furthermore, the lowest release level of MB observed at high pH was also consistent with the observed “closed” state of the gel (no GST) in the alkaline environment.

In contrast, levofloxacin (L) release studies showed a different trend (Figure 7, right). Although a GST was observed for all pH values, levofloxacin 24h release rates did not appear to mirror that GST trend. Indeed levofloxacin drug release rates increased with pH, suggesting that an alkaline environment was more favorable to L release. Again, this unexpected behavior may be due to the bulky nature of L compared to that for the MB more “linear” molecule, and/or an internal repulsion effect between the negatively charged L drug and the negatively charged physical crosslinks within the gel, as aforementioned. Thus, the pH release response behavior of levofloxacin loaded gel TMSL compared to that for MB loaded gel TMS+ differed greatly. This is likely due to the nature of the pH dependent physical interactions between the gel and the L drug molecule.

A broader conclusion to be made from these results is the following: pH sensitivity of borate based crosslinked physical reversible PVA release systems is likely affected by the nature of the loaded drug used. These results suggest that the physicochemical characteristics of the drug may play a key role in the pH response behavior of PVA borate based crosslinked gels (drug molecule size, drug pKa values and functional moieties present on the drug).

### Biocompatibility evaluation

Cytotoxicity results are shown in Figure 8. Crtl- and crtl+ are the non cytotoxic and cytotoxic samples respectively. TMS- and TMS+ are the gels with and without MB tested immediately after preparation (t = 0 h) or freeze-dried overnight (t = 24h). Results indicate good gel biocompatibility with a cell survival rate significantly above 70% deemed as the threshold value above which the material may be considered non-cytotoxic in accordance to the ISO10993-5 standard. These results are promising since they point out to the ability of our gels to be used in the context of wound dressings, tissue reparation or tissue regeneration systems.

**Figure 8.**
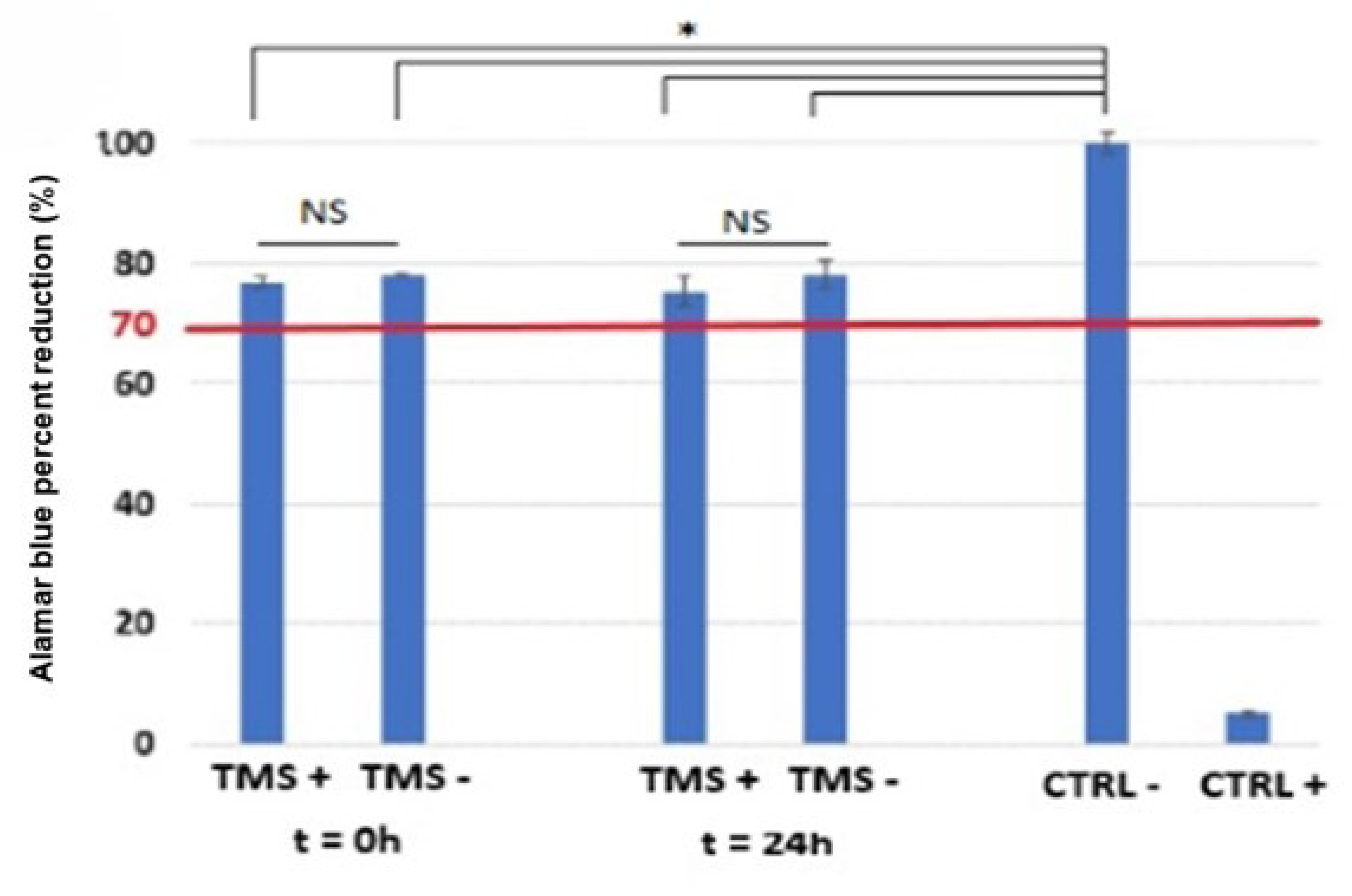
Alamar blue cytotoxicity assay for TMS gels.

Direct contacting test results (supporting information) are consistent with results above since they also indicate non-cytotoxicity for the gels (no cell growth disruption observed).

Furthermore, Live/Dead ® cell assay results (Figure 9) are consistent with non-cytotoxicity results above, since all cells remained alive (green) for short or long exposure times. These results are consistent with material non-cytotoxicity confirming overall good biocompatibility for our TMS systems.

**Figure 9.**
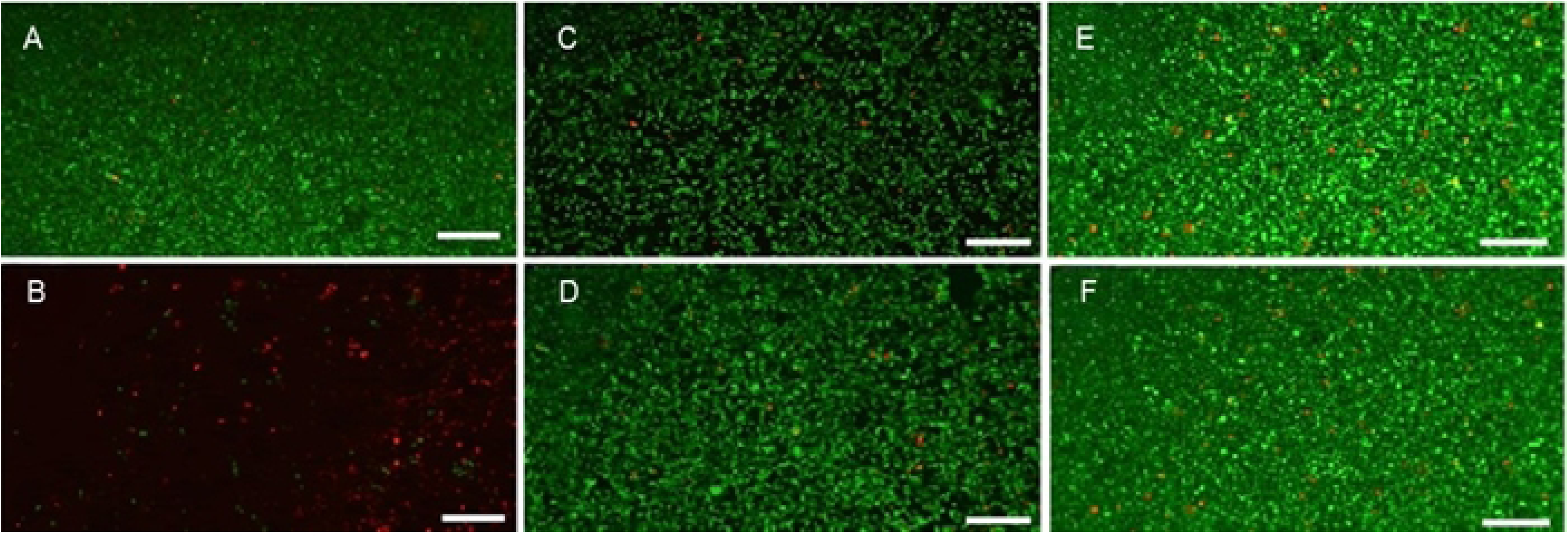
Cytotoxicity Live/Dead® assay. Fluorescence microscopy of L929 cells for negative (A) and positive cytotoxic control (B). (C) and (E) are the TMS+ gels (0 h) and (24 h), respectively. (D) and (F) are the TMS gels (0 h) and (24 h), respectively. Red and green indicate dead and alive cells, respectively. Scale bar represents 200 µm.

## Conclusion

We were able to demonstrate the feasibility of the elaboration of a cost effective PVA based reversible gel capable of releasing a drug in the presence of a specific biomarker of a chronic state. We also demonstrated that gel release profiles were dependent on the nature of the drug. Furthermore, we were able to confirm that these gels were biocompatible in the context of chronic wound care and/or tissue repair. These results are promising since they open the way to a new cost-effective elaboration strategy for ROS bio-responsive gels.

## Acknowledgments

The authors acknowledge Évreux Portes de Normandie (EPN) for financial support of the UMR CNRS 6270 PBS - BioMMAT team. This work was partially supported by Normandie Université (NU), the Région Normandie, the Centre National de la Recherche Scientifique (CNRS) and Université de Rouen Normandie (URN).

